# Non-Random Mis-Segregation of Human Chromosomes

**DOI:** 10.1101/278697

**Authors:** J. T. Worrall, N. Tamura, N. Shaikh, A. Mazzagatti, T. van Lingen, B. Bakker, D. C. J. Spierings, E. Vladimirou, F. Foijer, S. E. McClelland

## Abstract

Recurrent patterns of chromosomal changes (aneuploidy) are widespread in cancer. These patterns are mainly attributed to selection processes due to an assumption that human chromosomes carry equal chance of being mis-segregated into daughter cells when fidelity of cell division is compromised. Human chromosomes vary widely in size, gene density and other parameters that might generate bias in mis-segregation rates, however technological limitations have precluded a systematic and high throughput analysis of chromosome-specific aneuploidy. Here, using fluorescence *In-Situ* hybridization (FISH) imaging of specific centromeres coupled with high-throughput single cell analysis, as well as single-cell sequencing we show that human chromosome mis-segregation is non-random. Merotelic kinetochore attachment induced by nocodazole washout leads to elevated aneuploidy of a subset of chromosomes, and high rates of anaphase lagging of chromosomes 1 and 2. Mechanistically, we show that these chromosomes are prone to cohesion fatigue that results in anaphase lagging upon release from nocodazole or Eg5 inhibition. Our findings suggest that inherent properties of specific chromosomes can influence chromosome mis-segregation and aneuploidy, with implications for studies on aneuploidy in human disease.

## Introduction

Aneuploidy – deviation from a multiple of the haploid chromosome number - is the leading cause of spontaneous miscarriage and birth defects in humans (Nagaoka et al., 2012) and represents a key hallmark of cancer (Mitelman) where recurrent patterns of aneuploidy, or somatic copy number alteration (SCNA) are observed (Ben-David et al., 2016; Duijf et al., 2013; Faggioli et al., 2012; Mitelman; Nagaoka et al., 2012). SCNA patterns in cancer are a consequence of ‘chromosome alteration’ rate driven by chromosomal instability (CIN), which is associated with poor prognosis for cancer patients (Birkbak et al., 2011; Jamal-Hanjani et al., 2015; Laughney et al., 2015; Lee et al., 2011; McClelland, 2017) coupled to evolutionary selection. Human chromosomes vary widely in size, gene density, interphase nuclear territory and heterochromatin distribution (**Figure 1a, Table S1**). However, the question of whether these or additional characteristics generate bias in mis-segregation rates has not been answered to date since high-throughput methods to analyse chromosome-specific aneuploidy are lacking. The standard approach to measure aneuploidy, manual scoring of chromosome number using Fluorescence *In-Situ* Hybridization (FISH) of centromere-targeted probes (Burrell et al., 2013; Fenech, 2007) is low-throughput and subject to significant artefacts (Faggioli et al., 2012; Knouse et al., 2014; Valind et al., 2013; van den Bos et al., 2016), limiting the resolution of previous efforts to examine biased mis-segregation (Brown et al., 1983; Evans and Wise, 2011; Fauth et al., 1998; Hovhannisyan et al., 2016; Spence et al., 2006; Torosantucci et al., 2009; Xi et al., 1997). New technologies such as next generation sequencing-based methods (Bakker et al., 2016; van den Bos et al., 2016) are still expensive and technically challenging (Bakker et al., 2015). To resolve this we analysed individual chromosome aneuploidy rates in a high throughput manner using a novel platform and in the absence of fitness effects and selection. We used the ImageStream^X®^ cytometer to quantify FISH-marked centromeres in thousands of single cells, following pharmacological induction of merotelic attachments (kinetochores attached to microtubules emanating from both spindle poles). We show that human chromosome mis-segregation is non-random, and validate our findings using single cell sequencing. Observation of specific chromosomes during mitosis reveals that chromosomes 1 and 2 are highly prone to lagging at anaphase in multiple non-transformed cell lines. Lastly, we elucidate the molecular pathway underlying this bias, and show that chromosomes 1 and 2 are particularly to prone to cohesion fatigue resulting in mal-attached kinetochores and lagging at anaphase.

**Figure 1.**
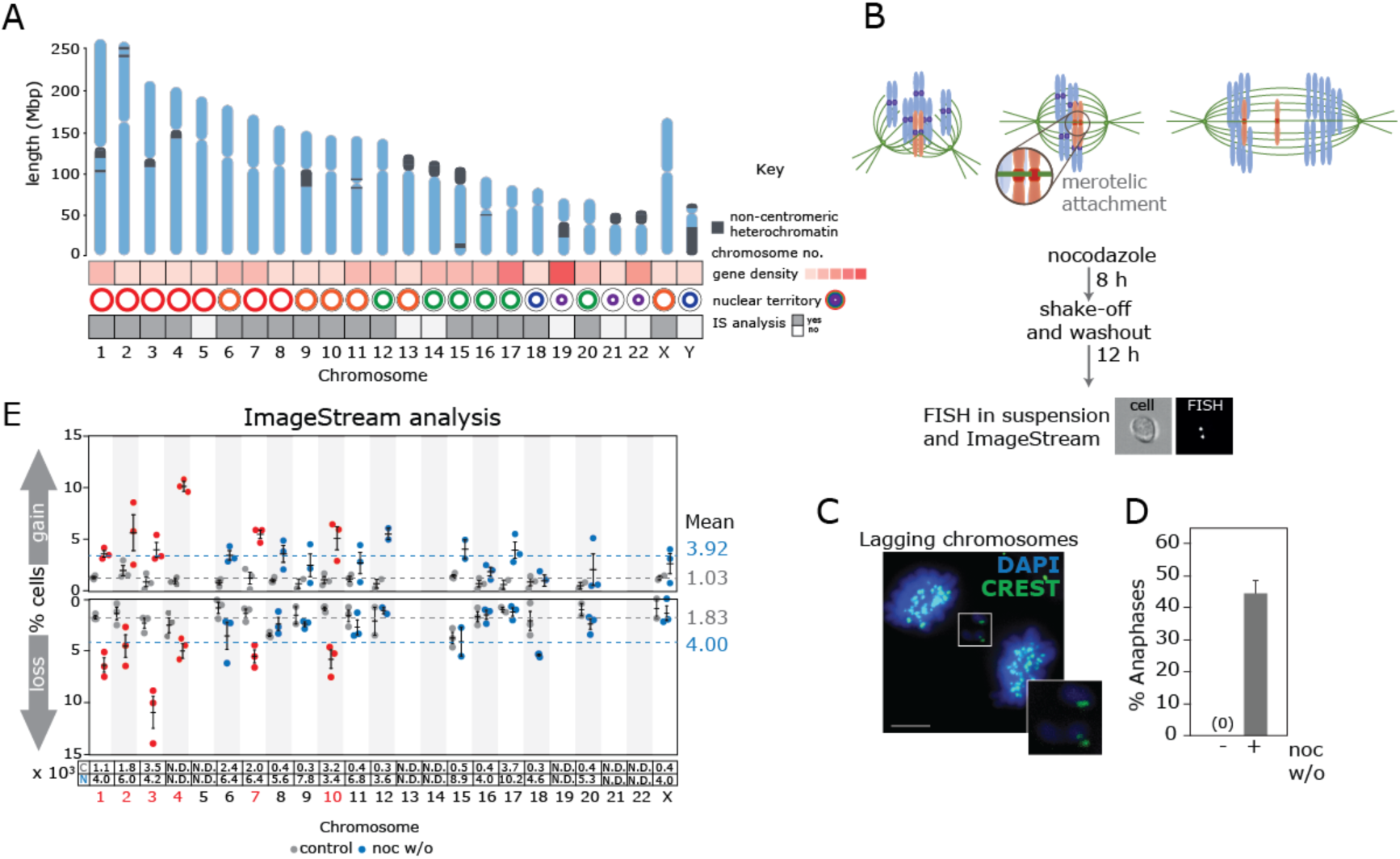
Chromosome mis-segregation induced by nocodazole washout leads to non-random aneuploidy in daughter cells. (**A**) Cartoon illustrating a selection of known chromosomal attributes (Cremer and Cremer, 2010). Gene density (no. genes divided by length of chromosome (Mbp)) was divided equally into 5 groups; 10.12-16.51; 16.52-22.89; 22.90-29.28; 29.29-35.67; 35.68-42.05. (**B**) Workflow of nocodazole washout and ImageStream analysis. (**C**) Immunofluorescence images of chromosome segregation errors in RPE1 anaphase cells as indicated following nocodazole washout. DNA is marked by DAPI (blue), centromeres are marked by CREST anti-sera (green). (**D**) Quantification of chromosome segregation errors in RPE1 anaphase cells following 8 h nocodazole followed by 1 h washout (mean and standard deviation of 2-3 experiments; 110 cells in total). (**E**) ImageStream analysis of RPE1 cells treated with nocodazole washout (light blue/red dots) compared to control (grey dots). Dots represent independent experiments and red dots and text mark chromosomes with mean nocodazole-induced aneuploidy rates above the mean for both gain and loss. Dashed lines indicate mean aneuploidy rates (grey:control; blue:nocodazole washout). Number of cells analysed per chromosome is indicated in lower box (x10^3^), (C=control; N=nocodazole washout).

## Results

### High throughput screening using the ImageStream^X®^ cytometer reveals non-random aneuploidy following induction of chromosome mis-segregation

Proposed drivers of cancer chromosomal instability include merotelic kinetochore attachments (Bakhoum et al., 2009a), replication stress (Bartkova et al., 2005; Bester et al., 2011; Burrell et al., 2013) and cohesion defects (Kim et al., 2016; Manning et al., 2014; Solomon et al., 2011). We examined the response of individual chromosomes to merotely in h-TERT-immortalised human retinal pigment epithelium cells (RPE1), using a nocodazole shake-off and washout strategy to induce merotelic attachment and chromosome mis-segregation (Cimini et al., 2002; Cimini et al., 2001; Zhang et al., 2015) (**Figure 1b,c**). To determine aneuploidy rates in the absence of selective pressure, we analysed cells 12 hours after release from nocodazole. Live cell imaging revealed that at this time-point cells have exited mitosis and proceeded through the cell cycle but without cell death or further division events that could influence population aneuploidy rates (**Figure S1a-c; Movie S1**) in agreement with previous reports that aneuploidy-mediated cell death does not occur in this timeframe (Li et al., 2010; Thompson and Compton, 2010). We also verified that the nocodazole treatment combined with mitotic shake-off and release does not impact cell viability (**Figure S1d-f**). We then performed high-throughput analysis of aneuploidy in daughter cells using the ImageStream^X®^ cytometer (hereafter ImageStream), previously employed to detect monosomy and trisomy in peripheral blood mononuclear cells with high accuracy (Minderman et al., 2012), to analyse individual chromosome aneuploidy frequencies at orders of magnitude higher cell number than previous approaches. We were able to perform this analysis for the majority of the 23 human chromosomes (**Figure 1a,e; Table S2**). As expected we observed an increase in overall aneuploidy following nocodazole washout. However, aneuploidy rates appeared varied between chromosomes, with chromosomes 1-4, 7 and 10 exhibiting mean aneuploidy rates above the average aneuploidy induced by nocodazole washout (**Figure 1e**).

### Single cell sequencing confirms ImageStream aneuploidy analysis

To validate ImageStream aneuploidy analysis we performed single cell sequencing (SCS) and aneuploidy detection using AneuFinder (Bakker et al., 2016) of control and nocodazole washout-treated RPE1 cells. Although cell numbers are orders of magnitude lower than the ImageStream analysis, SCS confirmed elevated aneuploidy for chromosomes 1, 2 and 3 (**Figure 2a,b**). A notable exception is chromosome 4, that was not observed as aneuploid using single cell sequencing and furthermore displayed higher gain than loss rates using ImageStream analysis, suggesting an artefact linked to the potential cross-reactivity of this probe (**Table S2**).

**Figure 2.**
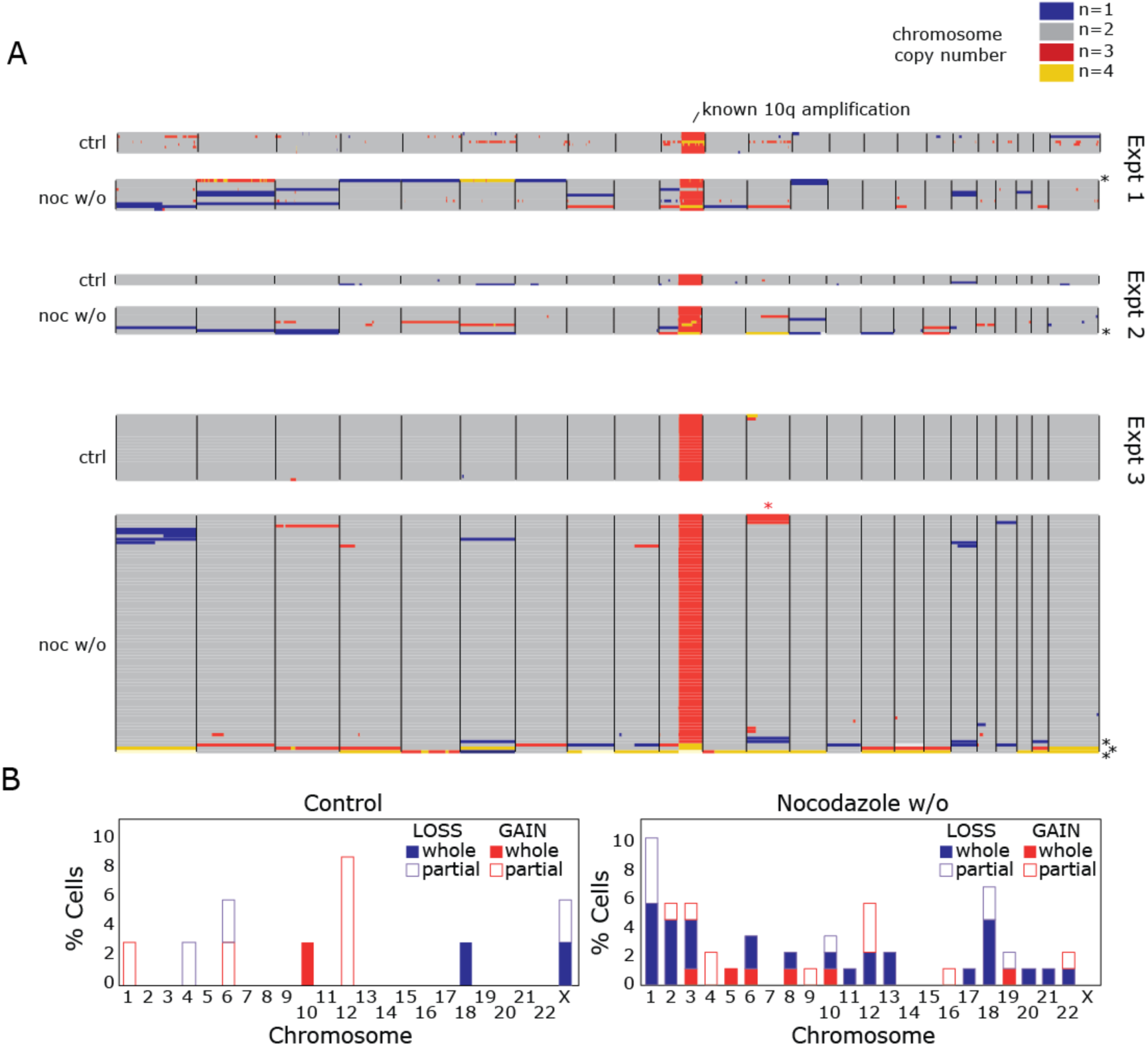
Single cell sequencing corroborates ImageStream aneuploidy patterns. (**A**) Genome-wide copy number profiles of control, and nocodazole washout treated RPE-1 cells from single cell sequencing data analysed using AneuFinder (Bakker et al., 2016) (three independent experiments; 34 control and 87 nocodazole w/o cells in total). Cells with more than 6 aneuploidies per cell were discarded from the analysis as this suggested a multipolar division (5 cells were removed under these criteria, black asterisks). Known subclonal gains of chromosome 12 were also excluded (red asterisks). Each row represents a single cell with chromosomes plotted as columns. Copy number states are depicted in different colours (see key). (**B**) % Cells exhibiting whole or partial aneuploidy events were collated from (**A**). Known amplification of chromosome 10q is caused by an unbalanced translocation to the X chromosome.

### Chromosomes 1 and 2 exhibit high rates of lagging at anaphase in multiple non-transformed cell types

We reasoned that chromosome-specific aneuploidy generated by nocodazole washout should be reflected in the propensity of specific chromosomes to undergo lagging during anaphase, though a strict relationship might not be expected since merotelically attached lagging chromosomes are often resolved to the correct daughter cell (Cimini et al., 2004; Thompson and Compton, 2011). To examine this, RPE1 cells were treated with nocodazole for 8 hours then released for 1 hour by which time approximately half the mitotic cell population had entered anaphase with approximately one in two cells displaying lagging chromosomes (**Figure S2a,b; Figure 3a,b**). We then performed FISH with specific centromere probes and determined the frequency of lagging of a panel of chromosomes, including those prone to aneuploidy following both ImageStream and SCS analysis. Strikingly, chromosomes 1 and 2 were found lagging in 56.4 ± 9 and 25.8 ± 2 % of anaphases with errors respectively (**Figure 3a-c**) suggesting these chromosomes comprised a high proportion of all lagging chromatids. To verify this we calculated lagging rates as a function of all lagging chromatids. Chromosomes 1 and 2 comprised 23.3 ± 7 % and 10.9 ± 3 % lagging chromatids respectively (**Figure 3d**) meaning that over a third of lagging chromatids following nocodazole washout are due to just two chromosomes. These observations were not due to non-specific signal since the number of specific centromere signals in each separating anaphase mass were scored to verify the correct total number of signals were present. Nocodazole washout also enriched lagging of chromosomes 1 and 2 in BJ cells, primary human umbilical endothelial cells (HUVEC) and h-TERT-immortalised fallopian epithelial cells (FNE1) (**Figure 3e-h, Figure S2c-j**), demonstrating that this effect is common to multiple non-transformed cell lines of different origins. We next used an alternative method to elevate chromosome mis-segregation: Small molecule inhibition of the Eg5 kinesin using monastrol or S-Trityl-L-Cysteine (STLC) prevents centrosome separation at prophase, leading to monopolar spindles. Upon drug washout spindles reform but promote merotelic attachment (Kapoor et al., 2000). Monastrol washout treatment induced similar total lagging chromosome rates and also strongly enriched lagging of chromosomes 1 and 2 (**Figure S2k-n**) demonstrating that sensitivity of chromosomes 1 and 2 was not due to nocodazole, or depolymerisation of microtubules (MTs) *per se*. Non-random chromosome-specific behaviour can thus be observed during error-prone mitosis, further supporting the conclusion that bias in aneuploidy rates can occur independently of selective pressure.

**Figure 3.**
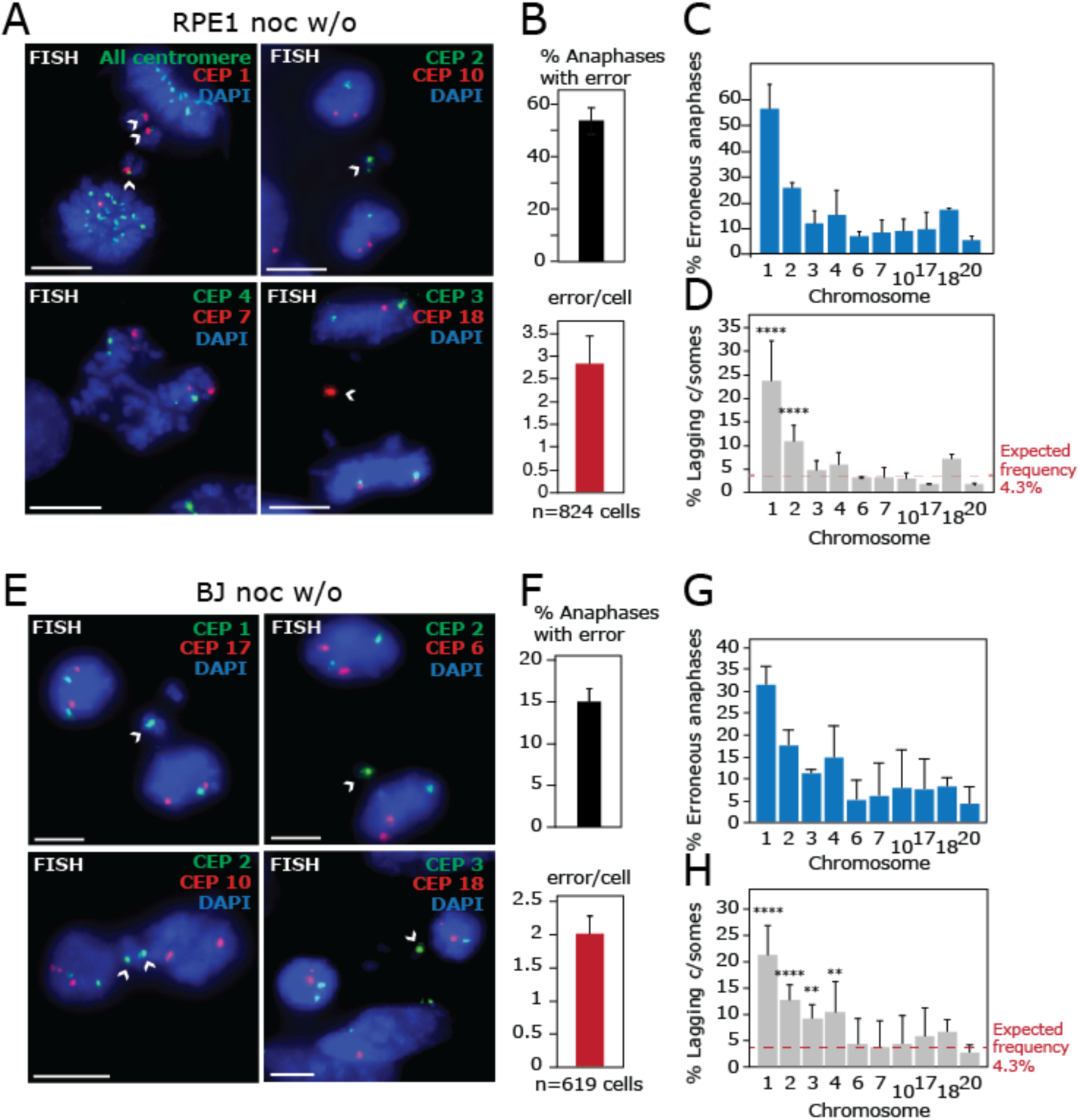
Chromosome 1 is highly prone to lagging at anaphase after nocodazole washout. (**A**) RPE1 cells were treated with 8 h nocodazole then released for 1 h before FISH was performed using all centromere, or specific centromere enumeration probes as indicated. (**B**) Segregation error rates and average number of lagging chromosomes per cell with error. Mean and standard deviation of six independent experiments is shown. (**C**) % Erroneous RPE1 anaphases (≥1 lagging chromosomes) exhibiting lagging of chromosomes indicated. (**D**) Specific chromosome lagging rates calculated as a function of all lagging chromatids. Total lagging chromatids were scored using DAPI-positive chromatid counting. Expected frequency is calculated using 1/23 assuming a random distribution among the 23 human chromosomes. (C) and (D) show mean and standard deviation of three independent experiments (except chr 17; two experiments), 149-251 cells, and 268-481 lagging chromosomes analysed per chromosome. (**E**) BJ cells were treated with 8 h nocodazole then released for 1 h before FISH was performed using specific centromere enumeration probes as indicated. (**F**) Segregation error rates and average number of lagging chromosomes per cell with error. Mean and standard deviation of four independent experiments is shown. (**G**) % Erroneous BJ anaphases (≥1 lagging chromosomes) exhibiting lagging of chromosomes indicated. (**H**) Quantification of % of lagging chromatids that are the chromosome indicated from erroneous anaphases. (G) and (H) show mean and standard deviation from three independent experiments, 84-157 cells, and 144-307 lagging chromosomes analysed per chromosome. P values in (D) and (H) were determined using a binomial test with Bonferroni multiple testing correction applied (see Methods) **<0.005, ****<0.00005. All scales bars 10 µM.

### Enrichment of chromosome 1 lagging is specific to treatments that interfere with normal bipolar spindle assembly

We next addressed the molecular mechanism underlying the sensitivity of chromosomes 1 and 2 to merotelic attachment following nocodazole or Eg5 inhibitor washout. To test whether an alternative method to induce mis-segregation lead to a similar phenomenon we treated cells with Reversine, a small molecule inhibitor of the mitotic checkpoint kinase Mps1 that promotes merotelic attachment and ablates the mitotic checkpoint (Santaguida et al., 2010). This treatment induced similar overall lagging chromosome rates however the bias for chromosome 1 lagging was removed (**Figure 4a-c**). This suggests that chromosome 1 lagging is promoted specifically by nocodazole or Eg5 inhibitor treatment and release.

**Figure 4.**
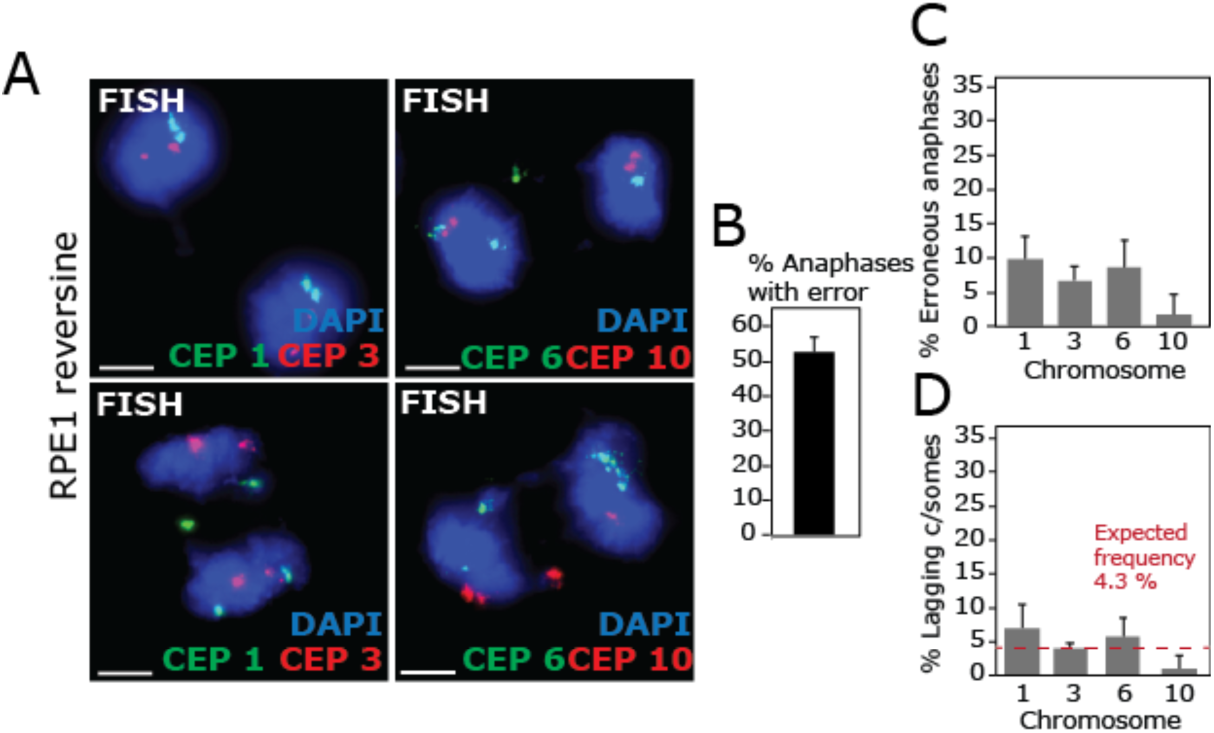
Enrichment of chromosome 1 lagging is specific to treatments that perturb bipolar spindle formation. (**A**) RPE1 cells were treated with 250 nM Reversine for 5 h to induce lagging chromosomes before FISH with centromeric probes as indicated. (**B**) % Anaphases with lagging chromosomes was quantified (n = 320 cells). (**C**) % Erroneous RPE1 anaphases (≥1 lagging chromosomes) exhibiting lagging of chromosomes indicated. (**D**) Quantification of % of lagging chromatids that are the chromosome indicated from erroneous anaphases. N = 251 erroneous anaphases in total in (C) and (D). All experiments show mean and standard deviation of three experiments. All scale bars 5 µm.

### Chromosomes 1 and 2 are susceptible to cohesion fatigue during mitotic delay

In addition to causing cells to pass through abnormal mitotic spindle geometry intermediates, both nocodazole and Eg5 inhibitor washout treatments are often coupled to a period of mitotic arrest, to elevate the number of mitotic cells available for analysis. A known consequence of delay in mitosis is gradual failure of the cohesive force holding sister chromatids together, ‘cohesion fatigue’, that can lead to premature sister chromatid separation (PSCS) (Daum et al., 2011; Manning et al., 2010; Nakajima et al., 2007; Stevens et al., 2011; van Harn et al., 2010) although underlying mechanisms remain unclear. Moreover, cohesion defects are known to promote chromosome lagging as shown by studies depleting the Retinoblastoma protein pRb (Manning et al., 2010) or STAG2, a component of the cohesin complex (Kleyman et al., 2014). To test whether cohesion fatigue was occurring during the 8-hour nocodazole treatment used herein, and whether this preferentially affected chromosomes 1 and 2, cells were treated with nocodazole for 8 hours followed by release into the proteasome inhibitor MG132 to delay anaphase but allow chromosome-MT attachments to form. Under these conditions cells displayed cohesion fatigue in the form of chromosomes ‘scattered’ from the metaphase plate (**Figure 5a,b**), albeit at lower rates than cells treated with MG132 alone for 8 h, in agreement with previous literature suggesting that dynamic MTs during the arrest period are required for maximal cohesion fatigue (Daum et al., 2011; Stevens et al., 2011). Chromosomes 1 and 2 were selectively prone to PSCS, with 46.1 ± 7 and 30.2 ± 10 % of cells with one or more unaligned chromosomes exhibiting PSCS of chromosomes 1 and 2 respectively, compared to 5.6 to 15.3 % for other chromosomes tested (**Figure 5c,d**). This suggested that cohesion fatigue might underlie the high rates of lagging of these chromosomes.

**Figure 5.**
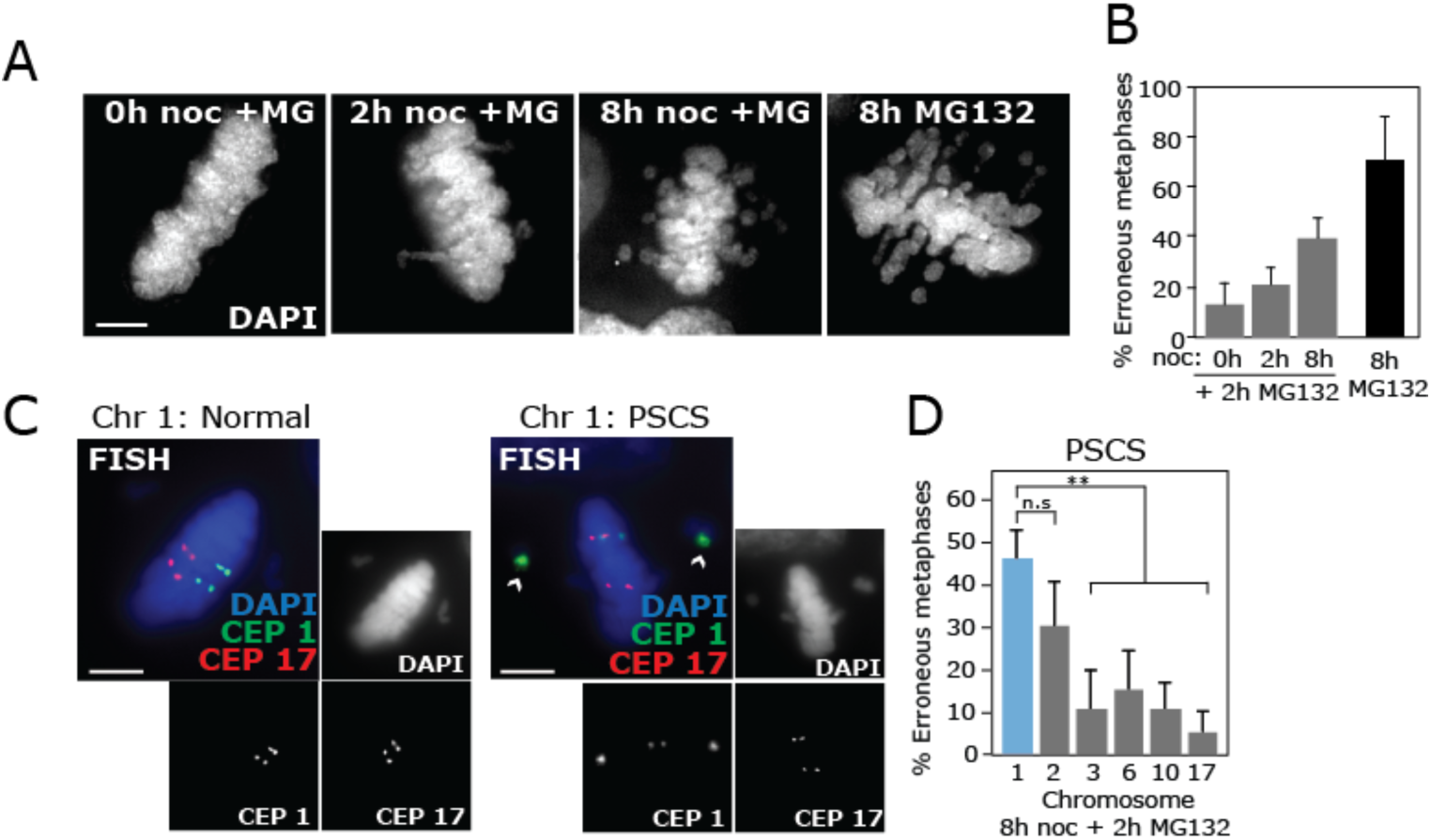
Mitotic-delay-mediated cohesion fatigue leads to PSCS of chromosomes 1 and 2. (**A, B**) RPE1 cells were treated with nocodazole as indicated then released into MG132 for 2 h, or treated with MG132 for 8 h, before scoring % cells with unaligned chromosomes (n = 300 cells in total). Significance was determined using one-way ANOVA with a post-hoc Tukey%s comparison test. (**C, D**) RPE1 cells were treated with 8 h nocodazole then 2 h MG132 before FISH with specific centromere enumeration probes (CEP) and quantification of PSCS (either one or both sister chromatids completely separated from metaphase plate) for each chromosome indicated. Erroneous metaphases (one or more unaligned chromosomes) exhibiting PSCS of a panel of chromosomes was quantified in (D) (108-141 cells in total per chromosome). All experiments show mean and standard deviation of three experiments. All scale bars 5 µm.

### Chromosome 1 lagging can be rescued by preventing cohesion fatigue

We then tested whether mis-segregation of chromosome 1 could be rescued by inhibiting cohesion fatigue. First, we depleted the negative regulator of cohesion, Wapl (Gandhi et al., 2006; Kueng et al., 2006) using RNA interference to enhance the stability of cohesion on DNA, previously shown to reduce PSCS and chromosome scattering at metaphase (Daum et al., 2011; de Lange et al., 2015; Lara-Gonzalez and Taylor, 2012; Stevens et al., 2011). This significantly reduced chromosome 1 mis-segregation following 8 hours nocodazole washout (**Figure 6a,b**). This treatment also reduced rates of mis-segregation globally (**Figure 6b**) likely reflecting both the contribution by chromosome 1 and a cumulative low-level effect upon other chromosomes. Next we reasoned that if cohesion fatigue induced by mitotic delay promotes lagging of chromosome 1, that reducing the length of mitotic arrest should rescue this effect. We therefore treated cells with a shorter period of nocodazole (2 hours) before washout. This reduced cohesion fatigue as measured by a decrease in scattered chromosomes and chromosome 1 PSCS of cells released into 2 h MG132 (**Figure 5b; Figure 6c,d**), and also reduced global rates of chromosome mis-segregation (**Figure 6e,f**). Importantly however, chromosome 1 lagging was preferentially rescued (**Figure 6f;** 4.50 ± 0.55 fold change compared to 2.74 ± 0.5 for global lagging rates). Reduced chromosome mis-segregation after 2 hours’ nocodazole washout was not due to incomplete MT depolymerisation or fewer cells affected by nocodazole, since mitotic cells displayed efficient loss of MTs after both 2 and 8 hours in nocodazole (**Figure S3a**), and live cell imaging of prometaphase cells released from nocodazole-induced mitotic arrest exhibited a similar reduction in segregation error rates between 8 and 2 hour treatments (**Figure 6g,h**). A similar reduction in lagging chromosome rates was also observed between 8 and 2 hours of Eg5 inhibition and release (**Figure S3b-d**). Taken together these data suggest that mitotic delay following spindle recovery from nocodazole or Eg5 inhibition leads to a deterioration of centromeric cohesion and a concomitant increase in chromosome lagging. This can be partially counteracted by increasing the stability of cohesion on DNA, or decreasing the length of mitotic arrest. Further, this phenomenon appears to particularly affect a subset of chromosomes with chromosomes 1 and 2 displaying both propensity to PSCS and high rates of lagging.

**Figure 6.**
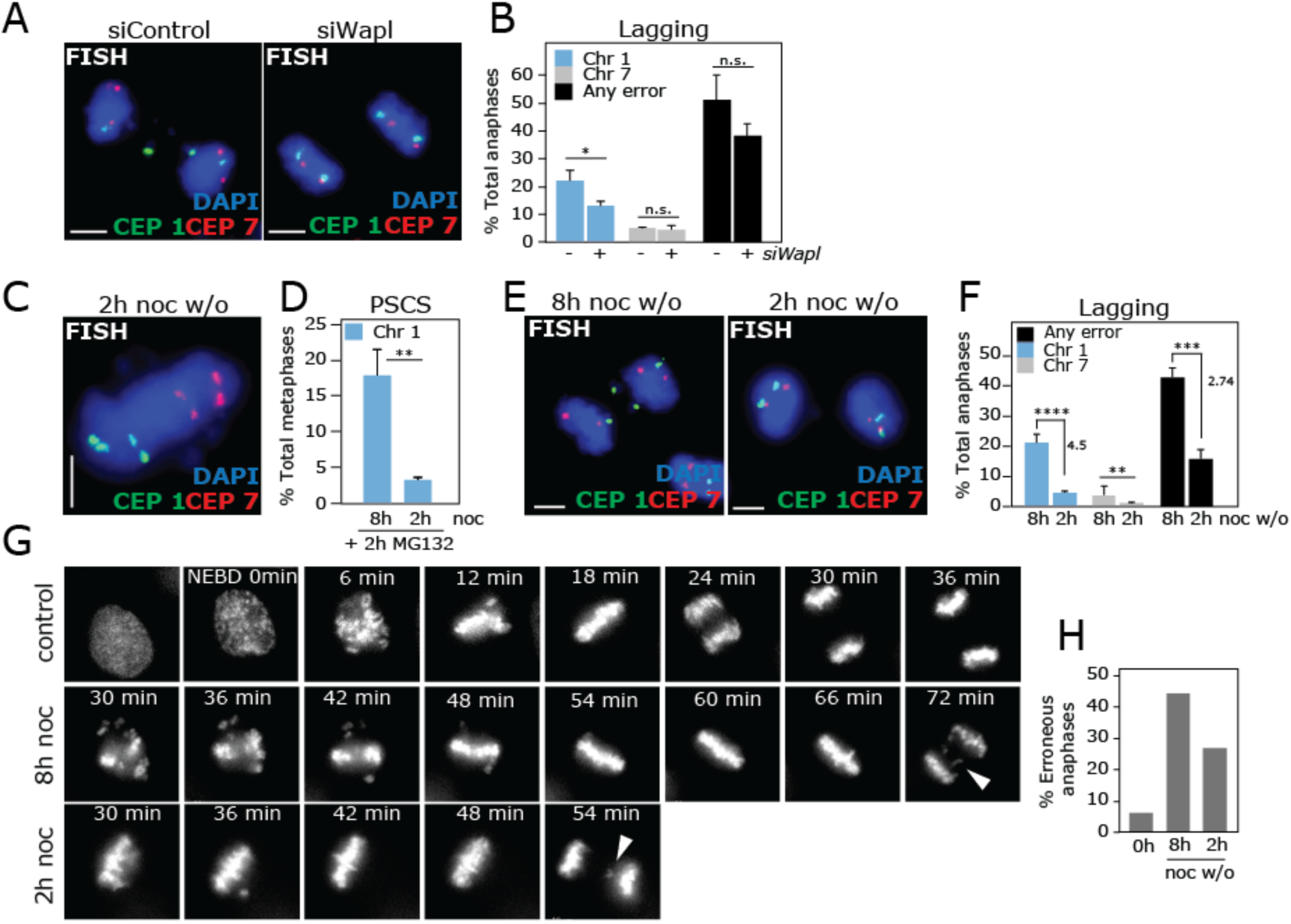
Chromosome 1 lagging can be rescued by reducing cohesion fatigue. (**A, B**) RPE1 cells were treated with non-targeting siRNA or siRNA targeted against Wapl for 39 h before 8 h nocodazole, washout for 1 h (48 h siRNA in total) then FISH. % Total anaphases with errors in any chromosome (n = 300 cells per condition), or specific chromosomes were analysed as indicated (B) (161 erroneous anaphases in total per condition). (**C, D**) RPE1 cells were treated with nocodazole for 2 h then released into MG132 for 2 h, before FISH and scoring of PSCS from erroneous metaphases (101 metaphases scored in total). (**E**) RPE1 anaphases after 2 or 8 h nocodazole treatment followed by 1 h release were prepared for FISH with centromere probes as indicated. (**F**) % Total anaphases with errors in any chromosome, chromosome 1, or chromosome 7 as indicated were calculated by first scoring errors from erroneous anaphases (n=43-46 (2 h) and 51-58 (8 h)) and converting error rates from erroneous anaphases to total anaphase rates (multiplying by % anaphase error rate (scored from 100 cells)/100). (**G**) Representative stills of movies of RPE1 cells stably expressing H2B-RFP, where filming began 30 minutes following washout from 2 or 8 h nocodazole treatment. (**H**) Quantification of anaphases with lagging chromosomes from live cell movies, summed from two independent experiments; 57, 57 and 95 cells from 0, 2 and 8 h nocodazole respectively. All experiments show mean and standard deviation of three experiments unless otherwise stated. P values were calculated using unpaired, two-tailed *t*-tests. All scale bars 5 µm.

## Discussion

In summary we demonstrate that compromising the fidelity of cell division can result in non-random mis-segregation, leading to differences in aneuploidy rates of individual human chromosomes. In this study we characterised non-random mis-segregation following induction of merotely using nocodazole washout. Alternative methods to induce aneuploidy could potentially affect different chromosomes, as a result of mechanistically distinct pathways to chromosome mis-segregation. Further investigation of such biases is likely to uncover novel aspects of chromosome-autonomous behaviour during faulty mitosis.

### Role of cohesion fatigue in promoting lagging of chromosomes 1 and 2

We find that cohesion fatigue during several hours’ delay at prometaphase particularly affects chromosomes 1 and 2, and is sufficient to promote a significant increase in their chromosome segregation error rate. Cohesion fatigue could elevate chromosome mis-segregation either due to effects on centromeric geometry or flexibility that might facilitate the formation of merotelic attachments (Sakuno et al., 2009). Alternatively, since multiple studies have demonstrated an intricate interplay between chromosome cohesion and the chromosomal passenger complex (CPC), responsible for error correction (reviewed in (Trivedi and Stukenberg, 2016); (Mirkovic and Oliveira, 2017)) it is possible that cohesion fatigue might prevent efficient correction of mal-attachments by improper regulation of the CPC.

### Cohesion fatigue induced by prometaphase delay also promotes chromosome mis-segregation in a global fashion

Although previous studies have suggested a requirement of dynamic MTs for cohesion fatigue (Daum et al., 2011; Stevens et al., 2011) our data show that a period of 8 hours in prometaphase, in the absence of a functional spindle, can predispose to loss of cohesion and chromosome scattering when cells are subsequently subjected to a brief period of MT pulling forces. This is in agreement with a recent study demonstrating increased sister chromatid separation after prolonging prometaphase using monastrol (Hengeveld et al., 2017) and indicates that cohesion fatigue may be more prevalent than previously appreciated during perturbations that induce a mitotic delay, even in the absence of dynamic MTs.

Moreover, in addition to promoting chromosome mis-segregation of specific chromosomes, we noted a marked dependency upon mitotic delay for high rates of global chromosome mis-segregation rates (**Figure 6f-h; Figure S3**). Subtle cohesion fatigue during mitotic delay thus appears to synergise with nocodazole or Eg5 inhibitor treatments to elevate chromosome mis-segregation, and may therefore play an unexpected role in generating segregation errors following these popular treatments to induce mis-segregation and aneuploidy.

### Features underlying bias in mis-segregation rates

Chromosomes 1 and 2 represent the largest chromosomes in humans (**Figure 1a**). Centromere repeat length does not scale with chromosome length in humans (**Table S1**) however longer chromosomes do not possess an obvious requirement for a ‘stronger’ centromere since drag produced by chromosomes is negligible in comparison to spindle forces (Civelekoglu-Scholey and Scholey, 2010; Nicklas, 1983). Nevertheless a correlation has been observed between chromosome size and levels of the inner centromeric protein CENP-A in human cells (Irvine et al., 2004) suggesting kinetochore size or function may vary between chromosomes. In this regard it is also interesting that chromosome 18, with the longest alpha satellite length (5.4 MB; **Table S1**) exhibited moderate but consistent effects in response to nocodazole washout both in terms of ImageStream aneuploidy loss rates and lagging chromosomes, despite falling short of statistical significance. This suggests that centromere size could also contribute to biased mis-segregation under certain conditions. Accordingly it has recently been shown in the Indian Muntjak that increased centromere size predisposes to merotelic attachment (H. Maiato, personal communication). It is possible that larger chromosomes may be prone to mis-segregation due to their tendency to occupy peripheral positions that might predispose to merotelic attachment (Cimini et al., 2004; Khodjakov and Rieder, 1996). Alternatively, it is possible that differences in centromeric composition underlie the sensitivity of chromosomes 1 and 2 to cohesion fatigue. Of note, large regions of pericentric heterochromatin have been identified at the q arms of chromosomes 1, 3, 4, 9, 16 and 19 (Atkin and Brito-Babapulle, 1981; Craig-Holmes and Shaw, 1971; Estandarte et al., 2016) (**Figure 1a**) although it is not clear whether the nature of chromosome 1 pericentric heterochromatin differs qualitatively, and how this might render chromosomes prone to cohesion fatigue and merotelic attachment.

### Potential role of non-random chromosome mis-segregation in development of cancer aneuploidy landscapes

Merotelic attachment and cohesion defects have both been proposed to contribute to cancer CIN (Bakhoum et al., 2009a; Bakhoum et al., 2009b; Ertych et al., 2014; Manning et al., 2014; Silkworth et al., 2009) therefore it is possible that aneuploidy-prone chromosomes identified herein may be subject to frequent mis-segregation and aneuploidy during tumourigenesis. However, confirming whether specific chromosomes are prone to mis-segregation during tumourigenesis is non-trivial. The bulk of available tumour genomic information lacks single cell resolution and is heavily shaped by evolutionary selection processes (Greaves and Maley, 2012; McGranahan and Swanton, 2017) that might obscure signatures of non-random mis-segregation. Nevertheless this phenomenon could influence early events during tumourigenesis. For example, lagging chromosomes can be subject to downstream DNA damage events such as breakage-fusion-bridge events and chromothripsis (Crasta et al., 2012; Janssen et al., 2011; Zhang et al., 2015) that could fuel subsequent structural aneuploidy events. In this regard it is interesting that chromosomes 1 and 2 are among the three chromosomes most frequently affected by copy number alteration in primary retinoblastomas (Kooi et al., 2016). Given links between dysfunction of the retinoblastoma protein pRB, cohesion defects and chromosome lagging (Manning et al., 2010; Manning et al., 2014), and the propensity for chromosomes 1 and 2 to lag under conditions of mal-attachment and cohesion fatigue in our hands, it is possible that non-random mis-segregation could act in concert with evolutionary selection to drive these recurrent SCNA patterns in retinoblastomas, and could potentially act more broadly across additional cancer types.

### Significance for interpretation of cancer genomes

Our study provides a framework for determining chromosome-level alteration rates that could improve the interpretation of SCNA landscapes from tumour genomes, for example, detection of ‘driver’ from ‘passenger’ SCNA events in cancer, since this requires the decoupling of mutation and selection rates (Beroukhim et al., 2010; Bignell et al., 2010). Moreover, if drivers of CIN that differ mechanistically affect distinct subsets of chromosomes, as discussed above, this leads to the possibility that analysis of non-random mis-segregation in cancer could allow us to decipher driver mechanisms, in a similar way to the recent discovery of cancer mutational signatures (Alexandrov et al., 2013). This also has implications for interpretation of mouse cancer models using specific drivers of CIN such as mitotic checkpoint knockouts, since the spectrum of errors may influence tumourigenesis and development (Ben-David et al., 2016).

## Author Contributions

J.W. N.T. N.S. A.M and S.M designed experiments. J.W. performed ImageStream analysis and FISH experiments in cells. N.T. performed live cell imaging, immunofluorescence and FISH experiments in cells. N.S. A.M. and T.vL. performed immunofluorescence and FISH experiments. D.S provided technical assistance with the single cell sequencing. B.B. analyzed the single-cell sequencing data. E.V. performed statistical analyses of lagging chromosome rates in Figure 3. F.F. provided resources and input on the project. S.M. conceived the study with input from J.W. and N.T., performed FISH experiments in cells and wrote the paper with contributions from all authors.

## Acknowledgments

We would like to thank Susana Godinho for the kind gift of BJ and RPE1 cells, and Tom Nightingale for HUVEC cells. We would also like to thank Elsa Logarinho, Patrick Meraldi, Daniele Fachinetti, Raquel Oliveira, Susana Godinho, Andrew McAinsh and Helder Maiato for helpful discussions. J.W. was funded by an MRC studentship, N.T. was funded by Barts and the London Charity (487/2133), N.S. was funded by PCRF, A.M. was funded by KKLF (KKL1073), TvL was funded by an ERASMUS studentship. F.F. and B.B were funded by the Dutch Cancer Society grant 2012-RUG-5549. We would like to thank the BCI Flow Facility and the Wellcome Trust for funding for the ImageStream (grant: 101604/Z/13/Z). Raw sequencing reads are available in the European Nucleotide Archive database (www.ebi.ac.uk/ena) under accession number TBD.”

---

**Figure S1.**
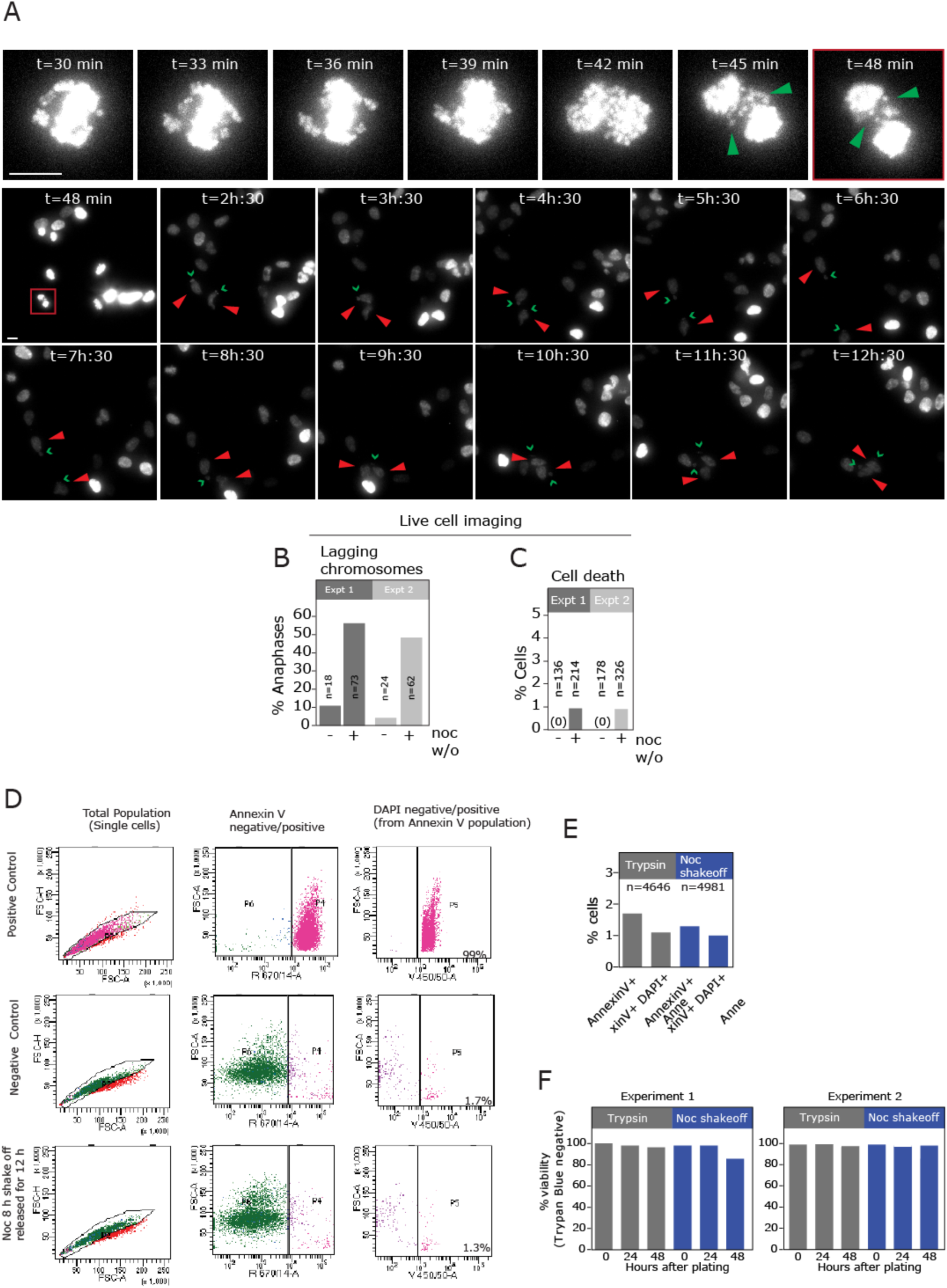
(relating to Figure 1). Aneuploidy-mediated cell death does not occur prior to ImageStream analysis. (**A-C**) RPE1 cells stably expressing H2B-RFP were filmed following release from 8 h nocodazole. Filming began 30 min after drug washout and cells were imaged every 3 min for 4 h, then every 15 min for a further 8 hours (12 hours% total filming). Segregation error rates (quantified from the first 4 hours of imaging; (B)) and cell death (C) rates were quantified from two independent movies. Stills from Supplementary Movie 1 are shown in (A). Green arrowhead indicates an anaphase cell with lagging chromosomes and chevrons indicate micronuclei formed from the lagging chromosomes. Red arrowheads mark daughter cells throughout the remainder of the movie. 39 daughter cells from mothers exhibiting lagging chromosomes could be followed for the full 12 h (cells frequently move ‘off screen’ during the subsequent hours). Scale bars 10 µM. (**D**) Flow cytometry plots of RPE1 cells subjected to Annexin V apoptosis assay. Positive control cells (top panels) were fixed prior to analysis to induce apoptosis and provide the gating for the untreated and nocodazole washout treated cells (middle and lower panels). (**E**) % Annexin V+ (early apoptotic) and Annexin V+, DAPI+ cells (late apoptosis) from (E). (**F**) Trypan blue cell viability assay of control, and nocodazole washout treated RPE1 cells treated with 8 h nocodazole then released for the times indicated.

**Figure S2.**
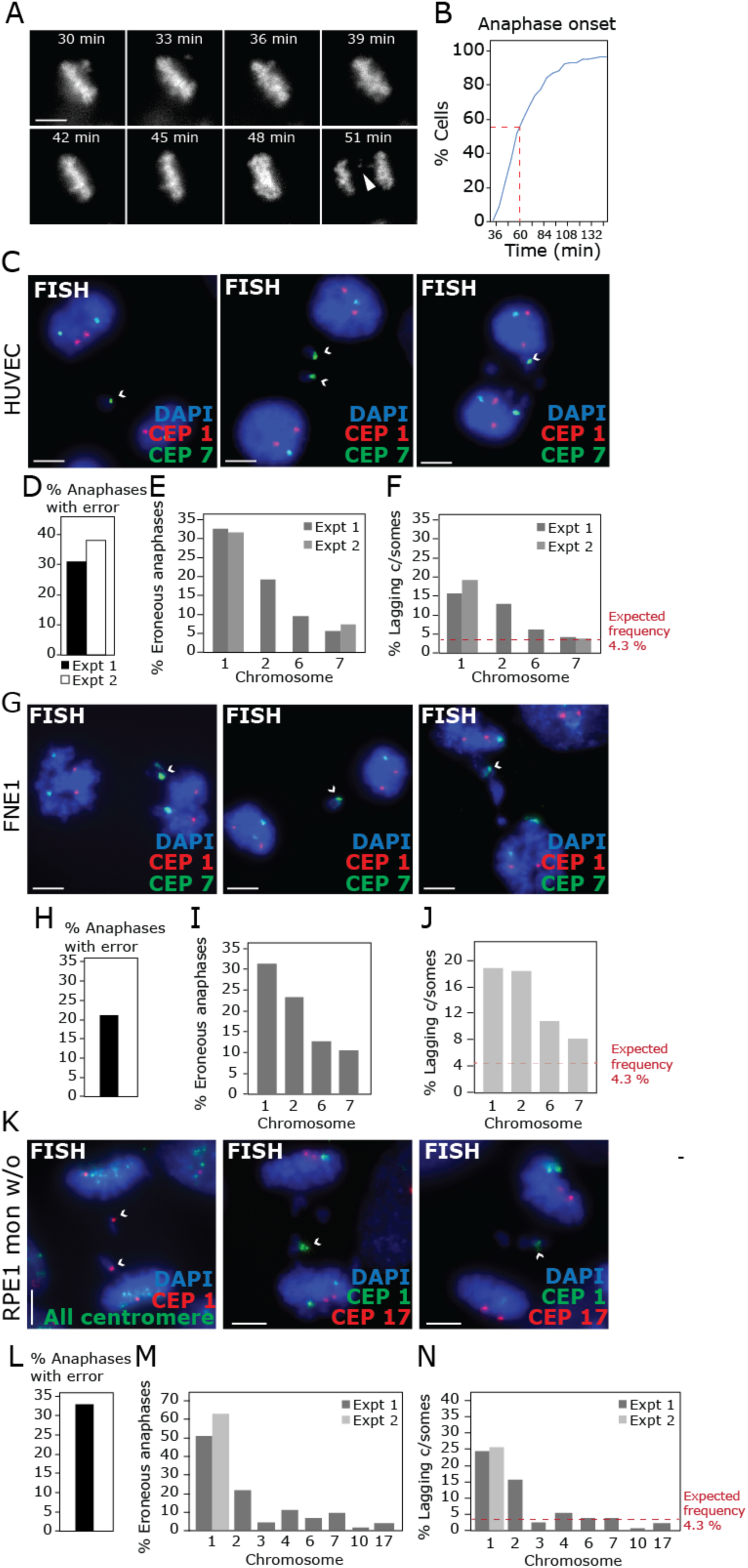
(relating to Figure 3). Chromosomes 1 and 2 are enriched in lagging chromosomes in multiple cell types. (**A**) Representative stills of movies of RPE1 cells stably expressing H2B-RFP, where filming began 30 minutes following washout from 8 h nocodazole treatment. (**B**) Cumulative frequency plot of timing of anaphase onset following release from nocodazole is shown from 95 cells in total from two independent experiments. (**C**) FISH image of chromosome 1 (green) and chromosome 7 (red) from HUVEC cells treated with nocodazole then released for 1 h. (**D**) % Anaphases with ≥1 lagging chromosome. (**E**) % Erroneous HUVEC anaphases (≥1 lagging chromosome) exhibiting lagging of chromosomes indicated. Results from 2 independent experiments are shown, 98 (CEP 1 and 7) and 52 (CEP 2 and 6) cells in total. (**F**) Quantification of % of lagging chromatids that are the chromosome indicated from erroneous anaphases (257 (CEP 1 and 2) and 147 (CEP 2 and 6) lagging chromosomes analysed in total. (**G)** FISH image of chromosome 1 (red) and all centromere (green) from FNE1 (fallopian tube epithelial) cells treated with nocodazole then released for 1 h. (**H**) % Anaphases with ≥1 lagging chromosome. (**I**) % Erroneous FNE1 anaphases (≥1 lagging chromosome) exhibiting lagging of chromosomes indicated. Results from 1 experiment are shown, 47-48 cells analysed per chromosome. (**J**) Quantification of % of lagging chromatids that are the chromosome indicated from erroneous anaphases. 85 (CEP 1 and 7) and 65 (CEP 2 and 6) lagging chromosomes analysed in total. (**K**) FISH image of chromosome 1 (red) and all centromere (green), or CEP 1 (green) and CEP 17 (red) from RPE1 cells treated with monastrol then released for 1.5 h. (**L**) % Anaphases with ≥1 lagging chromosome. (**M**) % Erroneous RPE1 anaphases (≥1 lagging chromosome) exhibiting lagging of chromosomes indicated. Results from 1-2 independent experiments are shown as indicated, 43-110 cells analysed per chromosome. (**N**) Quantification of % of lagging chromatids that are the chromosome indicated from erroneous anaphases. (77-299 lagging chromosomes analysed per chromosome). Images are projections of 5-15 0.2 µm z-slices. All scale bars 5 µm.

**Figure S3.**
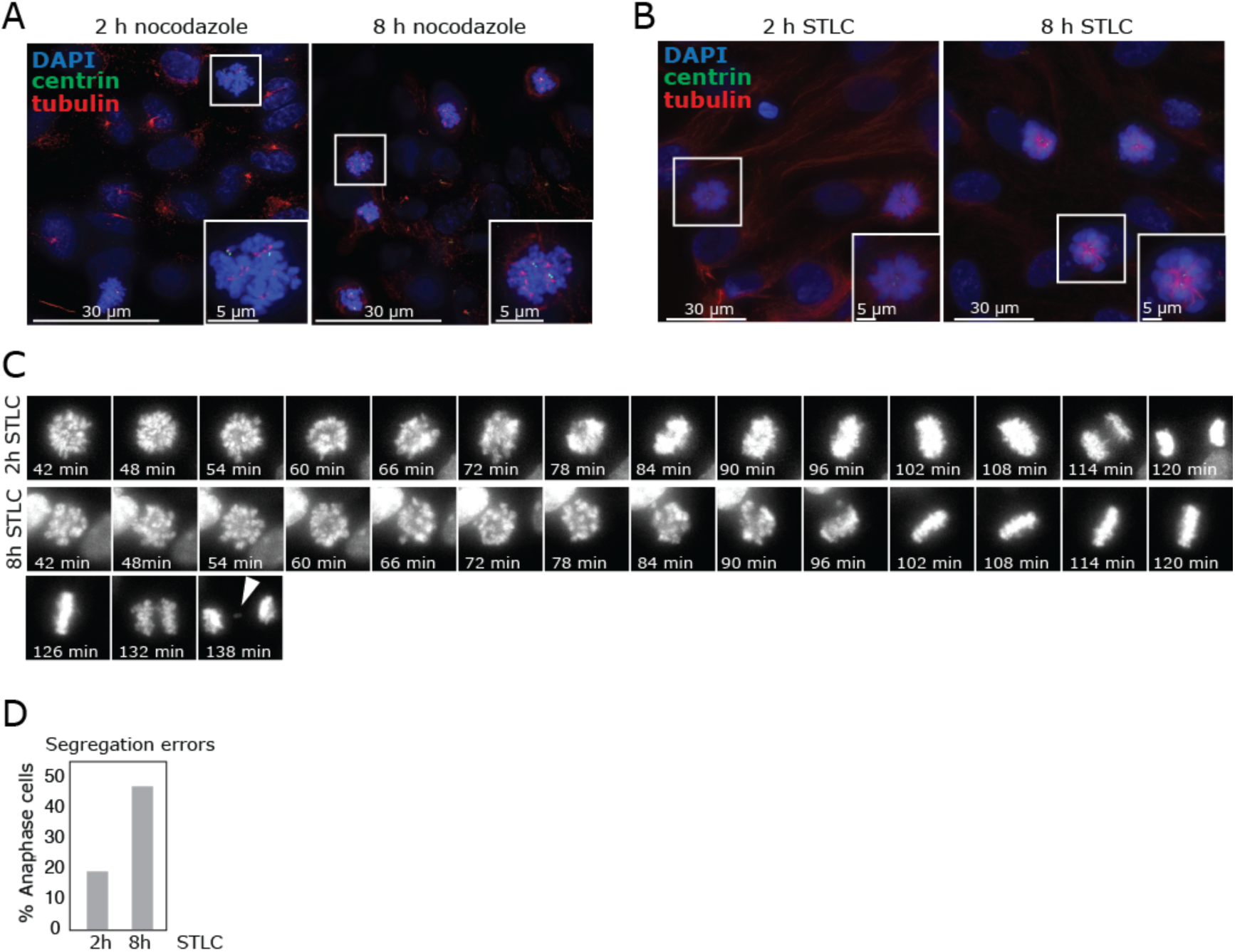
(relating to Figure 6). Shorter nocodazole or Eg5 inhibition treatment permits efficient spindle disruption but reduces chromosome mis-segregation rates. **(A,B)** RPE1 cells were fixed after 2 or 8 h nocodazole (A) or Eg5 inhibitor STLC (B) treatment (no washout) before staining with antibodies to beta-tubulin and centrin 3 to determine efficiency of MT depolymerisation. Images are projections. (**C**) RPE1 cells stably expressing H2B-RFP were filmed following release from 2 or 8 h STLC treatment. Filming began 30 minutes after release. (**D**) Quantification of anaphases with lagging chromosomes from live cell movies (78 (2 h STLC) and 104 (8 h STLC) cells in total from 2 independent experiments).

**Table S1.**
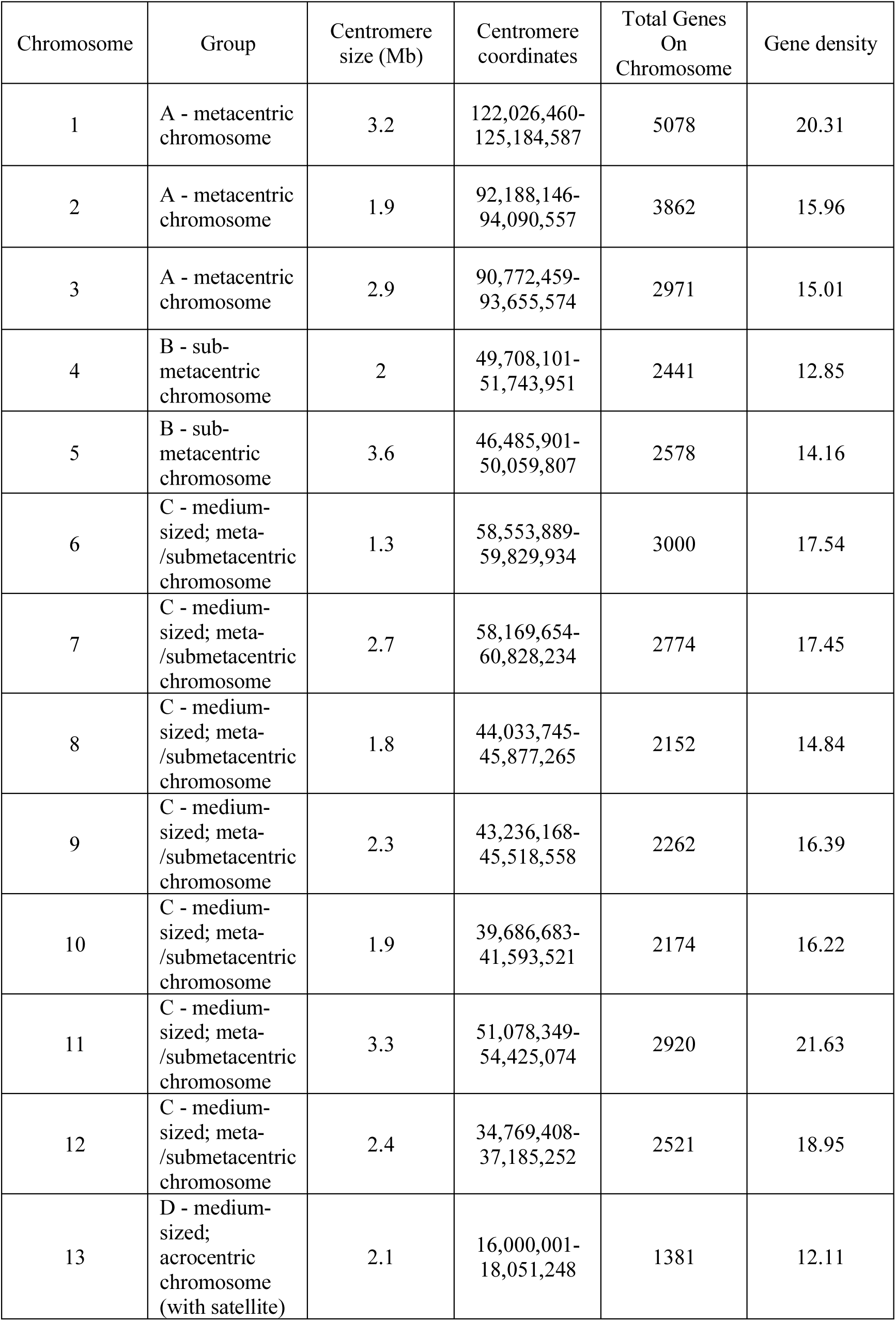
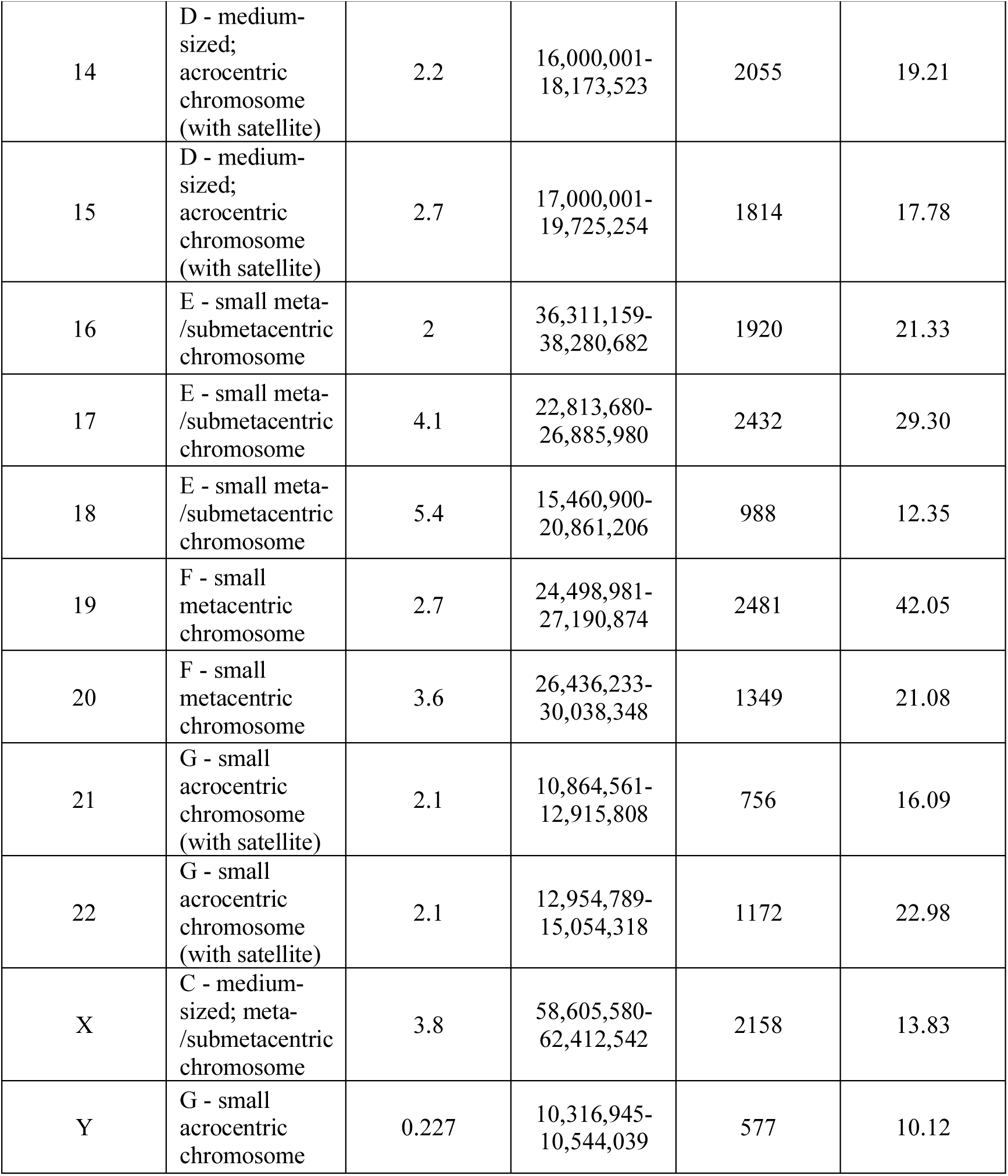
Chromosome characteristics.

| Chromosome | Group | Centromere size (Mb) | Centromere coordinates | Total Genes On Chromosome | Gene density |
| --- | --- | --- | --- | --- | --- |
| 1 | A - metacentric chromosome | 3.2 | 122,026,460-125,184,587 | 5078 | 20.31 |
| 2 | A - metacentric chromosome | 1.9 | 92,188,146-94,090,557 | 3862 | 15.96 |
| 3 | A - metacentric chromosome | 2.9 | 90,772,459-93,655,574 | 2971 | 15.01 |
| 4 | B - sub-metacentric chromosome | 2 | 49,708,101-51,743,951 | 2441 | 12.85 |
| 5 | B - sub-metacentric chromosome | 3.6 | 46,485,901-50,059,807 | 2578 | 14.16 |
| 6 | C - medium-sized; meta-/submetacentric chromosome | 1.3 | 58,553,889-59,829,934 | 3000 | 17.54 |
| 7 | C - medium-sized; meta-/submetacentric chromosome | 2.7 | 58,169,654-60,828,234 | 2774 | 17.45 |
| 8 | C - medium-sized; meta-/submetacentric chromosome | 1.8 | 44,033,745-45,877,265 | 2152 | 14.84 |
| 9 | C - medium-sized; meta-/submetacentric chromosome | 2.3 | 43,236,168-45,518,558 | 2262 | 16.39 |
| 10 | C - medium-sized; meta-/submetacentric chromosome | 1.9 | 39,686,683-41,593,521 | 2174 | 16.22 |
| 11 | C - medium-sized; meta-/submetacentric chromosome | 3.3 | 51,078,349-54,425,074 | 2920 | 21.63 |
| 12 | C - medium-sized; meta-/submetacentric chromosome | 2.4 | 34,769,408-37,185,252 | 2521 | 18.95 |
| 13 | D - medium-sized; acrocentric chromosome (with satellite) | 2.1 | 16,000,001-18,051,248 | 1381 | 12.11 |
| 14 | D - medium-sized; acrocentric chromosome (with satellite) | 2.2 | 16,000,001-18,173,523 | 2055 | 19.21 |
| 15 | D - medium-sized; acrocentric chromosome (with satellite) | 2.7 | 17,000,001-19,725,254 | 1814 | 17.78 |
| 16 | E - small meta-/submetacentric chromosome | 2 | 36,311,159-38,280,682 | 1920 | 21.33 |
| 17 | E - small meta-/submetacentric chromosome | 4.1 | 22,813,680-26,885,980 | 2432 | 29.30 |
| 18 | E - small meta-/submetacentric chromosome | 5.4 | 15,460,900-20,861,206 | 988 | 12.35 |
| 19 | F - small metacentric chromosome | 2.7 | 24,498,981-27,190,874 | 2481 | 42.05 |
| 20 | F - small metacentric chromosome | 3.6 | 26,436,233-30,038,348 | 1349 | 21.08 |
| 21 | G - small acrocentric chromosome (with satellite) | 2.1 | 10,864,561-12,915,808 | 756 | 16.09 |
| 22 | G - small acrocentric chromosome (with satellite) | 2.1 | 12,954,789-15,054,318 | 1172 | 22.98 |
| X | C - medium-sized; meta-/submetacentric chromosome | 3.8 | 58,605,580-62,412,542 | 2158 | 13.83 |
| Y | G - small acrocentric chromosome | 0.227 | 10,316,945-10,544,039 | 577 | 10.12 |

**Table S2.** Centromere probes.

| Chromosome | Probe | Chromosome Region | DNA Class | Notes |
| --- | --- | --- | --- | --- |
| 1 | CEP1 | 1q12 | Satellite III |  |
| 2 | CEP2 | 2p11.1-q11.1 | α-satellite | |
| 3 | CEP3 | 3p11.1-q11.1 | α-satellite | |
| 4 | CEP4 | 4p11.1-q11.1 | α-satellite | Shows occasional cross-reactivity with a group C chromosome (ref:Cytocell) |
| 5 | N/A |  |  | No specific centromeric probe available due to high sequence similarity between chromosomes 1, 5 and 19. Specific sub-telomere probe was tested but did not provide adequate signal to noise for ImageStream analysis. |
| 6 | CEP6 | 6p11.1-q11.1 | α-satellite | |
| 7 | CEP7 | 7p11.1-q11.1 | α-satellite | |
| 8 | CEP8 | 8p11.1-q11.1 | α-satellite | |
| 9 | CEP9 | 9q12 | Satellite III |  |
| 10 | CEP10 | 10p11.1-q11.1 | α-satellite | |
| 11 | CEP11 | 11p11.1-q11.1 | α-satellite | |
| 12 | CEP12 | 12p11.1-q11.1 | α-satellite | |
| 13 | N/A |  |  | No specific centromeric probe available due to high sequence similarity between other acrocentric chromosomes. Specific sub-telomere probe was tested but did not provide adequate signal to noise for ImageStream analysis. |
| 14 | N/A |  |  | No specific centromeric probe available due to high sequence similarity between other acrocentric chromosomes. Specific sub-telomere probe was tested but did not provide adequate signal to noise for ImageStream analysis. |
| 15 | CEP15 | 15p11.1-q11.1 | α-satellite | |
| 16 | CEP16 | 16p11.1-q11.1 | α-satellite | |
| 17 | CEP17 | 17p11.1-q11.1 | α-satellite | |
| 18 | CEP18 | 18p11.1-q11.1 | α-satellite | |
| 19 | N/A |  |  | No specific centromeric probe available due to high sequence similarity between chromosomes 1, 5 and 19. Specific sub-telomere probe was tested but did not provide adequate signal to noise for ImageStream analysis. |
| 20 | CEP20 | 20p11.1-q11.1 | α-satellite | Unequal intensity between homologous alleles observed. |
| 21 | N/A |  |  | No specific centromeric probe available due to high sequence similarity between other acrocentric chromosomes. Specific sub-telomere probe was tested but did not provide adequate signal to noise for ImageStream analysis. |
| 22 | N/A |  |  | No specific centromeric probe available due to high sequence similarity between other acrocentric chromosomes. Specific sub-telomere probe was tested but did not provide adequate signal to noise for ImageStream analysis. |
| X | CEPX | Xp11.1-q11.1 | α-satellite | |
| Y | N/A |  |  |  |

All CEP probes were from Cytocell.

## Movie S1

RPE1 cells stably expressing H2B-RFP were filmed following release from 8 h nocodazole treatment. Filming began 30 min after drug washout and cells were imaged every 3 min for 8 h, then every 15 min for a further 4 hours (12 hours% total filming). A Quicktime movie of one field is shown. Stills from this movie are shown in Figure S1.

## Materials and Methods

### Cell Culture and RNA interference

All cell lines were maintained at 37°C with 5% CO_2_. hTERT-RPE-1 cells (gift from S. Godhino) were cultured in DMEM Nutrient Mixture F12 Ham (Sigma); BJ cells (gift from S. Godhino) in DMEM high glucose (Sigma). Media for both was supplemented with 10% FBS and 100U Penicillin/Streptomycin. RPE1 and BJ cells were subjected to STR profiling to verify their identity in October 2017 using the cell line authentication service from Public Health England. HUVEC (Human umbilical cord vein endothelial cells, gift from T. Nightingale) cells were cultured in Huvec media (Medium 199, Gibco; 20% FBS; Endothelial Cell Growth Supplement, Sigma; 10U/ml Heparin, Sigma). FNE1 (University of Miami) cells were grown in FOMI media (University of Miami) supplemented with cholera toxin (Sigma). hTERT-RPE-1 H2B-RFP stable cell lines were generated after transfection with lentiviral construct H2B-RFP (Adgene26001, gift from S. Godhino). RNA Interference (Furlan-Magaril et al.) was achieved by transfection of cells for 48 h with 30 nM small interfering RNA (siRNA) from GE Healthcare using Lipofectamine RNAiMAX (Invitrogen) and Optimem (Gibco). siControl (D-001210-02) and siWAPL SMART pool (M-026287-01) (Dharmacon). Cells were treated with drugs at the following concentrations: MG132 (Sigma) 10 µM; Nocodazole (Sigma) 100 ng/ml; S-Trityl-L-Cysteine (STLC) (Sigma) 10 µM; Release from nocodazole, or STLC, was achieved by washing cells with media three times, then incubating in fresh media for 60 or 90 minutes, respectively.

### Apoptosis assay

Cells were re-plated after either only trypsinisation or after 8 h nocodazole treatment followed by mitotic shake-off. After 12 hours, cells were collected and then stained with Annexin V Alexafluor 647 antibody (ThermoFisher Scientific A23204) and DAPI, fixed in formaldehyde and analysed by FACS.

### Immunofluorescence

Cells grown on glass slides or coverslips were fixed with a PTEMF solution comprised of 0.2% Triton X-100, 0.02 M PIPES (pH 6.8), 0.01 M EGTA, 1 mM MgCl_2_ and 4% formaldehyde. After blocking with 3% BSA, cells were incubated with primary antibodies according to suppliers% instructions: Beta-tubulin (Abcam ab6046), Centrin 3 (Abcam ab54531), CenpA (Abcam ab13939), CREST (Antibodies Incorporated, 15-234-0001), Aurora B (Cambridge Bioscience, A300-431A-T), Phosph-Hist H2a.X (Millipore, 05-636). Secondary antibodies used were goat anti-mouse AlexaFluor 488 (A11017, Invitrogen), goat anti-rabbit AF594 (A11012, Invitrogen), and goat anti-human AF647 (109-606-088-JIR, Stratech). DNA was stained with DAPI (Roche) and coverslips mounted in Vectashield (Vector H-1000, Vector Laboratories).

### Fluorescence *In Situ* Hybridisation (FISH)

Cells were grown on glass slides, fixed in methanol/acetic acid, then put through an ethanol dehydration series. Cells were incubated overnight at 37°C with specific centromere enumeration probes (CEP) or chromosome paints (all from Cytocell) or pan-centromere probes (Cambio Ltd), then washed the following day with 0.25x SSC at 72°C followed by 2x SSC, 0.05% Tween. When measuring cohesion fatigue, Premature Sister Chromatid Separation (PSCS) was defined as where either one, or both the two centromere signals of one pair of sister chromatids were completely separated from the metaphase plate.

### FISH-In suspension

For ImageStream analysis, FISH was performed in suspension: Cells in log-phase growth were treated with 100 ng/ml nocodazole for eight hours and released following mitotic shake-off into fresh medium for 12 hours before analysis. Cells from all experimental conditions were harvested, as previously described, and fixed by adding freshly-prepared 3:1 methanol-glacial acetic acid drop-wise to a pellet of PBS-washed cells. For hybridisation, cells were washed with 1x PBS with 3% BSA twice for five minutes, pelleted, and resuspended in 0.05% Tween20 and 2x Saline-sodium Citrate (Okosun et al.) in PBS. 1 x10^6^ cells from this suspension were pelleted and the supernatant removed by pipetting. Cells were then resuspended in 40µL of complete hybridisation mixture containing 28 µL hybridisation buffer, 10µL nuclease-free H_2_O and 2 µL CEP probe. Denaturing and probe hybridisation were performed in a thermocycler under the following conditions: 80 °C (five minutes), 42 °C (9 to 16 hours) and an optional storage step of 4 °C. Following hybridisation, 200 µL of 2x SSC was added to each reaction mixture. Cells were pelleted and resuspended in 50 to 100 µL of 1x PBS before analysis (optional: DAPI, 1 µg/mL).

### Microscopy

Images were acquired using an Olympus DeltaVision RT microscope (Applied Precision, LLC) equipped with a Coolsnap HQ camera. Three-dimensional image stacks were acquired in 0.2 µm steps, using an Olympus ×100 or ×60 1.4 numerical aperture UPlanSApo oil immersion objective. Deconvolution of image stacks and quantitative measurements was performed with SoftWorx Explorer (Applied Precision, LLC). For live cell imaging, H2B-mRFP-labelled cells were grown and imaged in 4 well imaging dish (Greiner bio-one). 20 µm *z*-stacks (ten images) were acquired using an Olympus ×40 1.3 numerical aperture UPlanSApo oil immersion objective every 3 min for 8 h using a DeltaVision microscope in a temperature and CO_2_-controlled chamber. Analysis was performed using Softworx Explorer. To observe cell death after nocodazole washout, cells were imaged every 3 min for the first 4 h and then every 15 min for another 8 h.

### ImageStream cytometry analysis

All samples were analysed on the ImageStream cytometer by excitation with the blue laser with a power of 100 mW at a ‘high’ flow speed. Data obtained by the ImageStream were analysed in IDEAS 6.2 (Merck Millipore). Samples for each chromosome and experimental condition were obtained separately and contained within a single data file. For each sample a minimum of 500, and a maximum of 40,000, cells were analysed. Raw data files were opened in the IDEAS software package and the built-in compensation matrix applied. This correction is necessary to remove fluorescent noise introduced from the spatial alignment between channels, the flow speed, camera background normalisation and the level of brightfield gain. During acquisition, the EDF element was used to increase the focus range from 4 µm to 16 µm, allowing close to 100% of cells to be focused. Single cells are distinguished from cell aggregates by low area and high aspect ratio. The gating of single cells was manually verified by visual observation of brightfield images in the selected region. Plotting the Gradient root mean squared (RMS) value of the brightfield channel allowed only cells that were in-focus to be analysed. In-focus cells have a high Gradient RMS value. For some samples, where the hybridisation efficiency was less, a further gate was applied to select for only cells in the sample above a threshold of probe signal intensity. This was achieved by plotting the total intensity of fluorescence in each cell, versus the Raw Max Pixel intensity within the cell. Cells with hybridised probe have an average total fluorescence, and a high Raw Max Pixel intensity. Single, in-focus, hybridised cells were then analysed for the chromosomal content of a particular chromosome by applying a ‘spot mask’ and ‘spot counting’ feature to the centromere probe signals for each image. The masking parameters were determined on user-defined variables: the radius of the spot and the spot-to-background ratio (STBR). The STBR is the spot pixel value divided by the background fluorescence of the bright detail image. The spot mask therefore denotes a region that is of appropriate area to be considered a centromeric signal, and the boundary at which the signal diminishes. Where the radius value is x, this suggests that the denoted area of a single spot should have a minimum value of 2x+1 pixels. Regions that satisfy the spot mask criteria in single cells are enumerated by the spot-counting wizard. For the wizard to accurately determine chromosome ploidy, truth populations were denoted for both 2n-1 and 2n+1 cells for a minimum of 25 images. The wizard then compiles the common features for over 100 elements and assigns each image a spot count.

The images obtained of CEP spots are 2D projections of 3D images, to encompass the entire volume of the nucleus. If a cell is aligned so that the two centromere signals are in the same plane, they sometimes appear as a single focus, because they overlap following image projection. To correct for this, CEP signal intensity was plotted as a histogram from the original spot count data which correlates with the amount of probe hybridised, rather than the spot count. Disomic cells had a medium (M) intensity of hybridisation signal intensity, representing two spots. Cells with one spot that had lost a chromosome will fall below the value represented by two standard deviations above the mean fluorescent intensity; cells that had gained a chromosome will fall above two standard deviations of the mean of the hybridisation signal intensity. Events that are classified as one spot by the software usually fell into the medium range for intensity in the majority of cases. This suggests that, for the reasons stated above, they are disomic cells with aberrant ploidy-spot relationship. Cells designated as one spot that fell outside the 2 standard deviation window were deemed to be true monosomies. Cells designated as 2n+1 by the spot-counting wizard were manually verified by visual inspection of each image and correlating it with the 2 standard deviation cut-off above the mean diploid fluorescence intensity.

### Single cell Sequencing

Samples from control and experimentally-induced aneuploid cells were sorted by FACS prior to single-cell sequencing analysis using AneuFinder as previously reported(Bakker et al., 2016). Briefly, sequence reads are determined as non-overlapping bins with an average length of 1 Mb, a GC correction is applied, and binned sequences are analysed using a Hidden Markov model to determine the most likely copy number states. To negate the inherent sample variation introduced by sequencing single cells, a stringent quality control step was included that uses multivariate clustering to exclude libraries of insufficient quality. Chromosome copy number is plotted as a genome-wide state with clustering of cells based on the similarity of copy number profiles. Raw sequencing reads are available in the European Nucleotide Archive database (www.ebi.ac.uk/ena) under accession number TBD.”

### Statistical analyses

Unpaired *t*-test or one-way ANOVA with post-hoc Tukey’s comparison were used to test for levels of significance using either Excel or Prism (GraphPad). Asterisks have been used to denote the significance value between experimental conditions adhering to the following nomenclature: p<0.05 (*); p<0.01 (**); p<0.001 (***); p<0.0001 (****).

To check whether specific chromosomes occurred in lagging error more often we used a binomial test. If chromosomes lag equally, then a lagging chromosome is expected to have a given identity in *p* = 1/23 = 4.3% of cases, and the number of observed is distributed as *y* ∼ Binomial(*p,n*), where *n* is the number of lagging chromosomes in total. We used Bonferroni multiple testing correction when applying this test across all observed chromosomes.

